# Modeling of ATP Transport in an Axon: Effects of Spontaneous Neuron Firing and Mitochondrial Transfer via Tunneling Nanotubes

**DOI:** 10.1101/2025.01.14.633089

**Authors:** Andrey V. Kuznetsov

## Abstract

While electrical activity in neurons has been extensively studied, the transport and distribution of adenosine triphosphate (ATP), the primary cellular energy carrier, remain less understood, particularly in relation to metabolic processes in axons. ATP is primarily generated in mitochondria and consumed at synapses, the primary sites of energy demand. Even in healthy axons, approximately half of synaptic boutons lack stationary mitochondria, raising questions about ATP transport between boutons with and without mitochondrial ATP production. This study addresses two key questions: the role of spontaneous neuronal firing in maintaining ATP levels during periods of low energy demand and the ability of a single bouton with a donated mitochondrion to supply ATP to neighboring boutons lacking mitochondria. Using computational simulations, the study examines ATP transport under various firing patterns and mitochondrial distributions, incorporating factors such as quiescent periods, duty cycles, and ATP diffusivity. Spontaneous neuronal firing stabilizes ATP levels during periods of low energy demand, preventing reactive oxygen species (ROS) release from mitochondria. Simulations reveal that in neurons damaged by neurodegeneration, a single bouton containing a donated mitochondrion can support ATP levels in multiple empty boutons. However, as the number of empty boutons increases, ATP concentration declines, potentially falling below the critical threshold required for synaptic transmission.

**Nomenclature:** *a*_0_ kinetic constant characterizing the rate of ATP consumption in a bouton

*A*_*c*_ cross-sectional area of the axon

*C* ATP concentration per unit length of the axon

*C*_0_ typical value of ATP concentration per unit length of the axon

*C*_min_ minimum ATP concentration required to sustain synaptic transmission

*duty* duty cycle

*D* ATP diffusivity in the cytosol

*f* frequency at which neuron fires during the active phase

*i*_*active*_ number of action potentials that occur during the active phase

*i*_*total*_ total number of action potentials that propagate down the axon during the active phase plus the number of action potentials that were missed during the quiescent phase

*L* distance between boutons

*m* ATP production rate per unit length of a mitochondrion

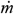 ATP production rate per unit mass of tissue

*N*_*empty*_ number of boutons in the CV that lack stationary mitochondria

*t* time

*x* position along the axon

*x*_*m*_ half-length of a mitochondrion

**Greek symbols:** *γ* percentage of tissue volume occupied by mitochondria

*δ* half the width of an axonal varicosity

*ϖ* mass of an individual mitochondrion

Λ_*homeostatic*_ portion of energy expended on homeostatic maintenance of a bouton

## 1. Introduction

While the electrical functioning of neurons has received significant attention [1,2], much less focus has been given to mass transport in neurons, particularly in relation to their metabolic processes. An axon is a slender, elongated structure extending from a neuron’s cell body, responsible for transmitting signals to connected cells. Signal transmission within the axon is electrical [3]. However, when the signal reaches a synapse, it transitions from electrical to chemical transmission. When activated, the synapse releases chemical messengers called neurotransmitters across a small gap, the synaptic cleft, which bind to receptors on the receiving cell [4]. Signal transmission is an energy-intensive process [5].

Adenosine triphosphate (ATP), a small molecule, serves as the primary energy carrier in cells, providing readily accessible chemical energy to power most cellular processes [6]. ATP breaks down into adenosine diphosphate (ADP) and a phosphate group, releasing energy in the process. Mitochondria (Fig. 1a) are organelles primarily responsible for cell energy production. Due to the high efficiency of glucose oxidation, mitochondria serve as the main source of cellular energy. The majority of ATP is generated from ADP through oxidative phosphorylation, a process that occurs within the mitochondria [7,8].

**Fig. 1.**
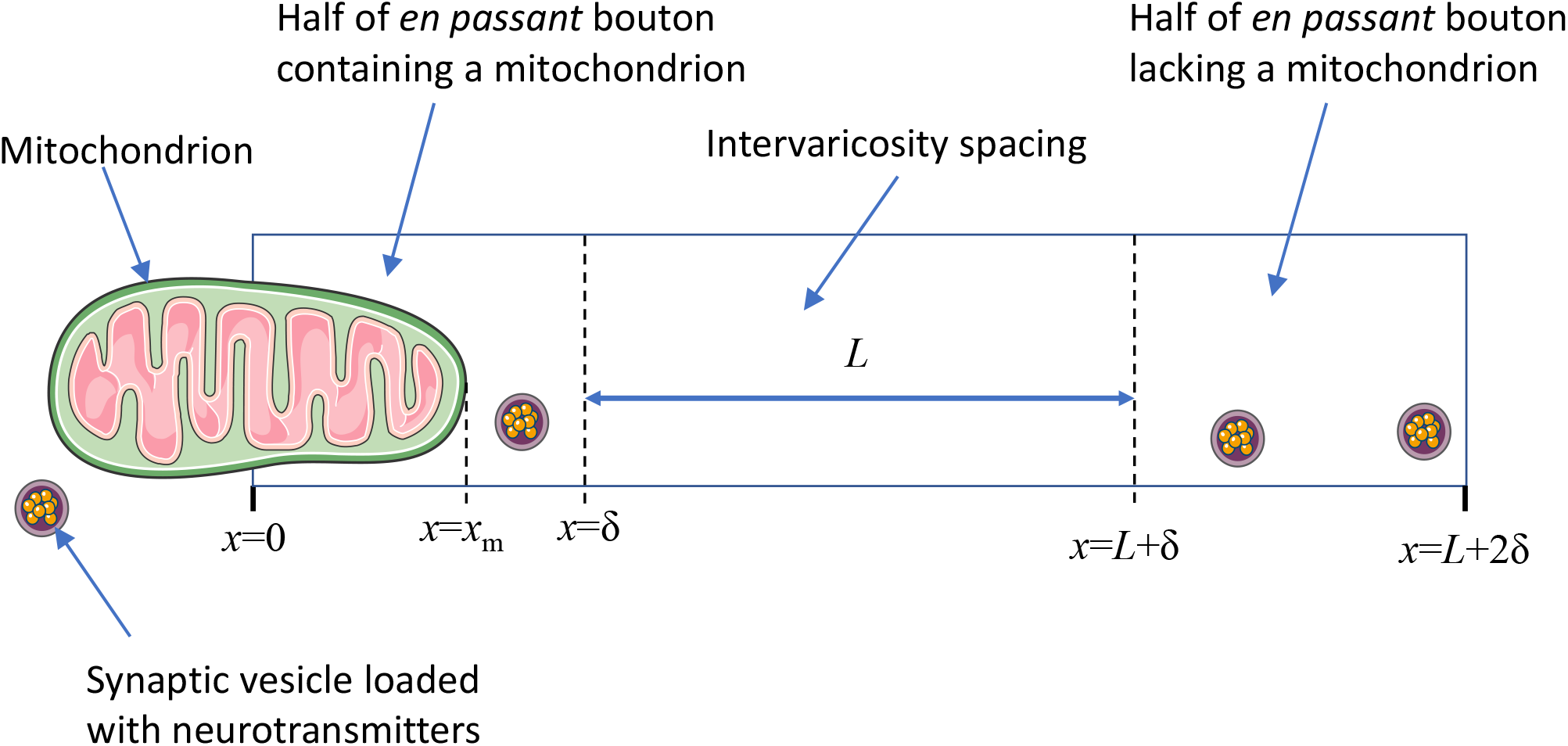
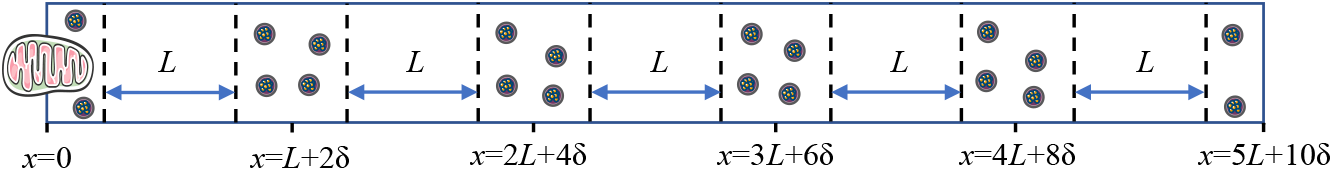
A schematic diagram illustrating a single periodic unit CV, which contains half of a mitochondrion, with the other half belonging to a neighboring periodic CV. The width of the axonal varicosity is denoted by 2 *δ*, and *L* represents the distance between consecutive varicosities. (a) Scenario showing a stationary mitochondrion present at every alternate varicosity. (b) Scenario showing a stationary mitochondrion present in every sixth varicosity, leaving five empty varicosities between those containing a mitochondrion. *Figure generated with the aid of servier medical art, licensed under a creative commons attribution 3*.*0 generic license*. http://Smart.servier.com.

Neurons communicate with other cells by releasing neurotransmitters at specialized structures known as synapses. Synapses release neurotransmitters in response to an arriving action potential. When situated along the axon shaft, these synapses appear as small varicosities and are known as en passant boutons [9] (hereafter referred to as boutons), see Fig. 1a. From this point forward, the terms synapses, boutons, and varicosities will be used interchangeably. Synapses are the primary sites of ATP consumption within the brain [5]. Refs. [10-13] indicate that less than half of en passant boutons contain stationary mitochondria, with the reason for this uneven distribution still unknown.

This paper examines two key questions about ATP transport in neurons, specifically addressing how ATP is transported between boutons with and without mitochondria. The first question examines how spontaneous neuronal firing during periods of low ATP consumption may protect neurons from toxic conditions in mitochondria. The traditional view of neural firing is that it functions primarily as a communication mechanism. However, many aspects of neural firing are not well understood. Many neural systems exhibit spontaneous activity, for example, during sleep, without any apparent external triggers. Ref. [14] proposed that this spontaneous firing plays a crucial metabolic role, acting as a protective mechanism to prevent the release of reactive oxygen species (ROS) from mitochondria, which can occur under conditions of low ATP consumption and production. ROS can be particularly damaging to neurons, as they cannot divide and must function for the entire lifetime. Therefore, one of the goals of this paper is to investigate the role of spontaneous neuronal firing in regulating ATP levels during periods of low energy demand in boutons with mitochondria and those without.

In 2004, Gerdes and colleagues [15] discovered tunneling nanotubes (TNTs), long membranous protrusions enabling cell-to-cell communication [16,17]. Mathematical models of the transport of intracellular components in TNTs were developed in refs. [18-20]. Recently, ref. [21] demonstrated that TNTs enable mitochondrial transfer from microglia to neurons, potentially mitigating mitochondrial dysfunction. However, given the fragility of TNTs and the limited number of donated mitochondria, questions remain about the effectiveness of this mechanism and its long-term impact on neuronal health [22]. TNTs remain stable for minutes to hours, enabling short-term cytoplasmic exchange between the connected cells [23]. Given the limited number of healthy mitochondria that can be donated to neurons during this time, the second question examined in this study is: how many empty boutons can a bouton with a single donated healthy mitochondrion supply with ATP while maintaining ATP concentrations in these boutons above the critical threshold?

To address these two questions, this study employs a computational approach to simulate ATP dynamics in axons under various firing patterns. The axon is divided into multiple periodic control volumes (CVs), each comprising one bouton with a mitochondrion and one or more boutons without mitochondria. By incorporating factors such as quiescent periods, duty cycles, and ATP diffusivity, the model provides insights into how neuronal firing and mitochondrial distribution influence ATP concentrations in axonal segments. The findings illuminate the metabolic implications of spontaneous firing and mitochondrial transfer, offering insights into mechanisms that support neuronal health and resilience.

## 2. Materials and models

The model focuses on a single dependent variable: the linear ATP concentration, denoted by *C*, which is defined as the number of ATP molecules per unit length of the axon. A summary of the model parameters is provided in Table 1. Equations given below are extensions of those developed in refs. [24,25].

**Table 1.**
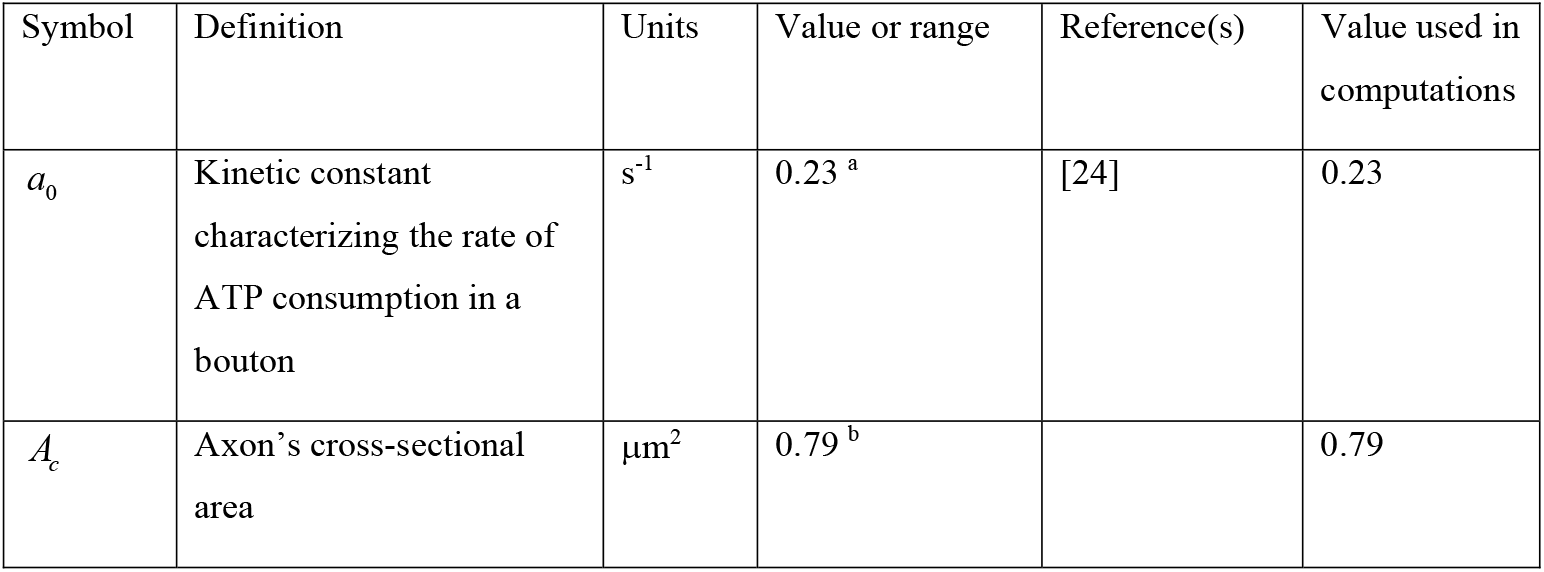
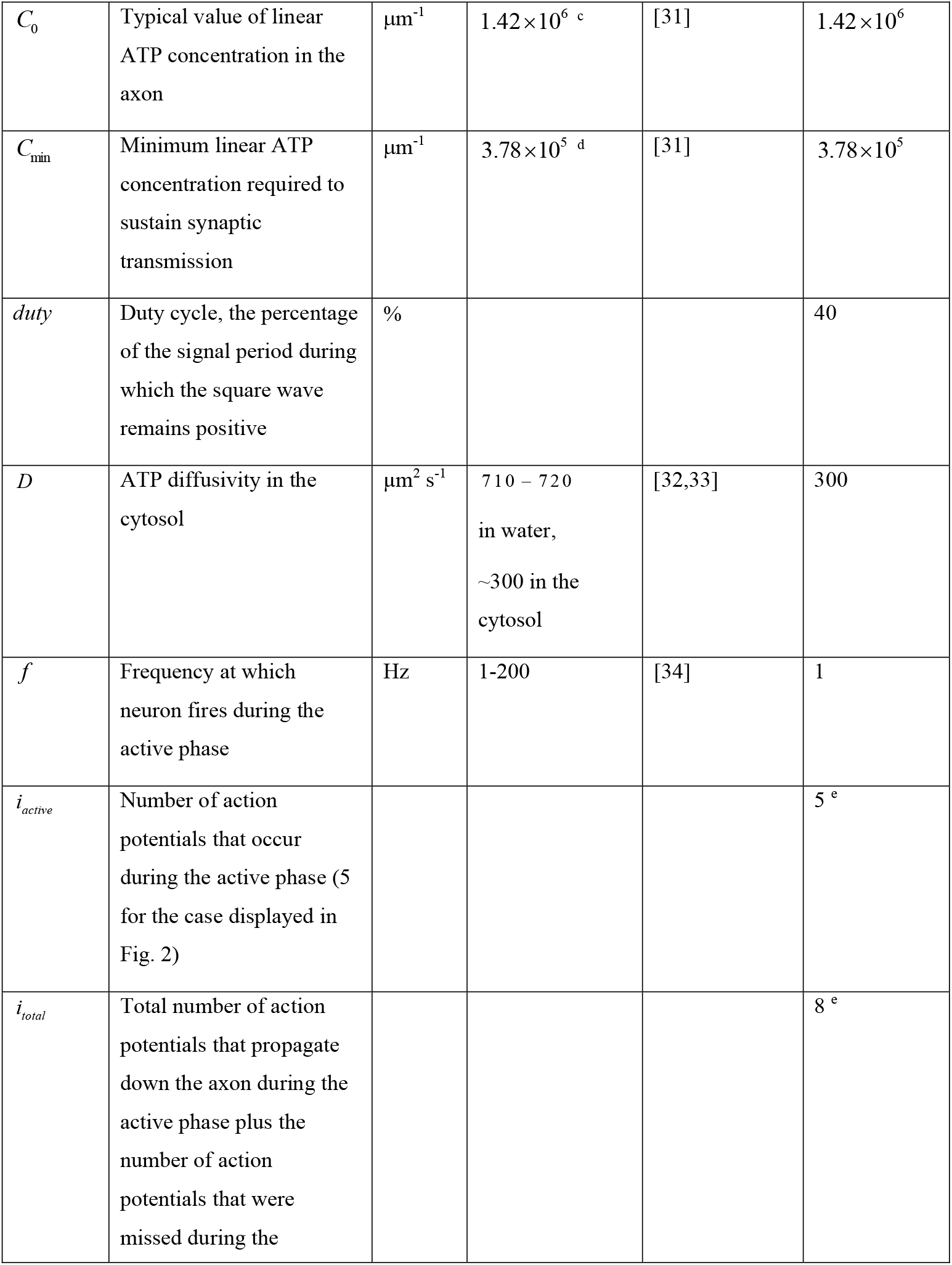
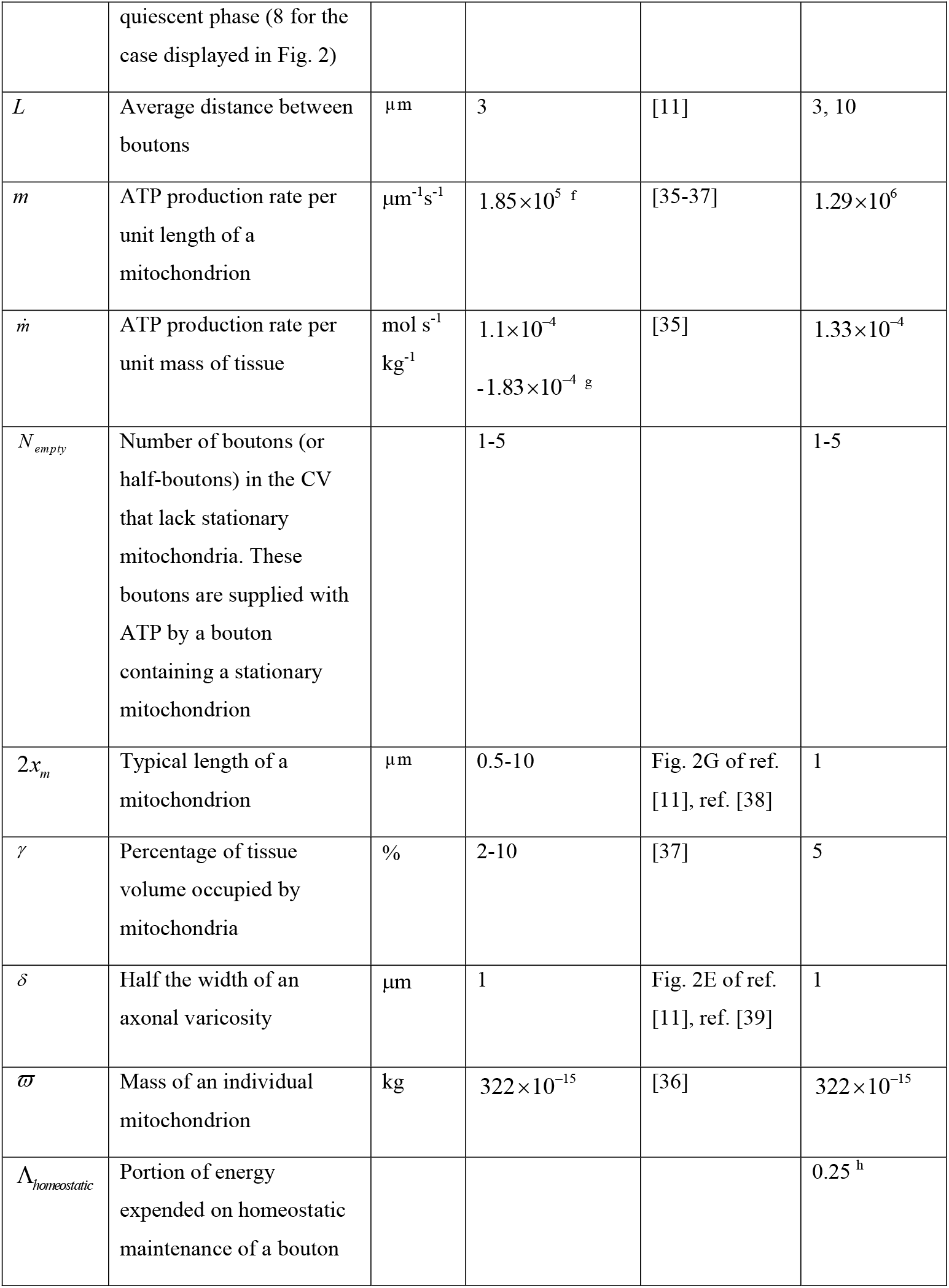
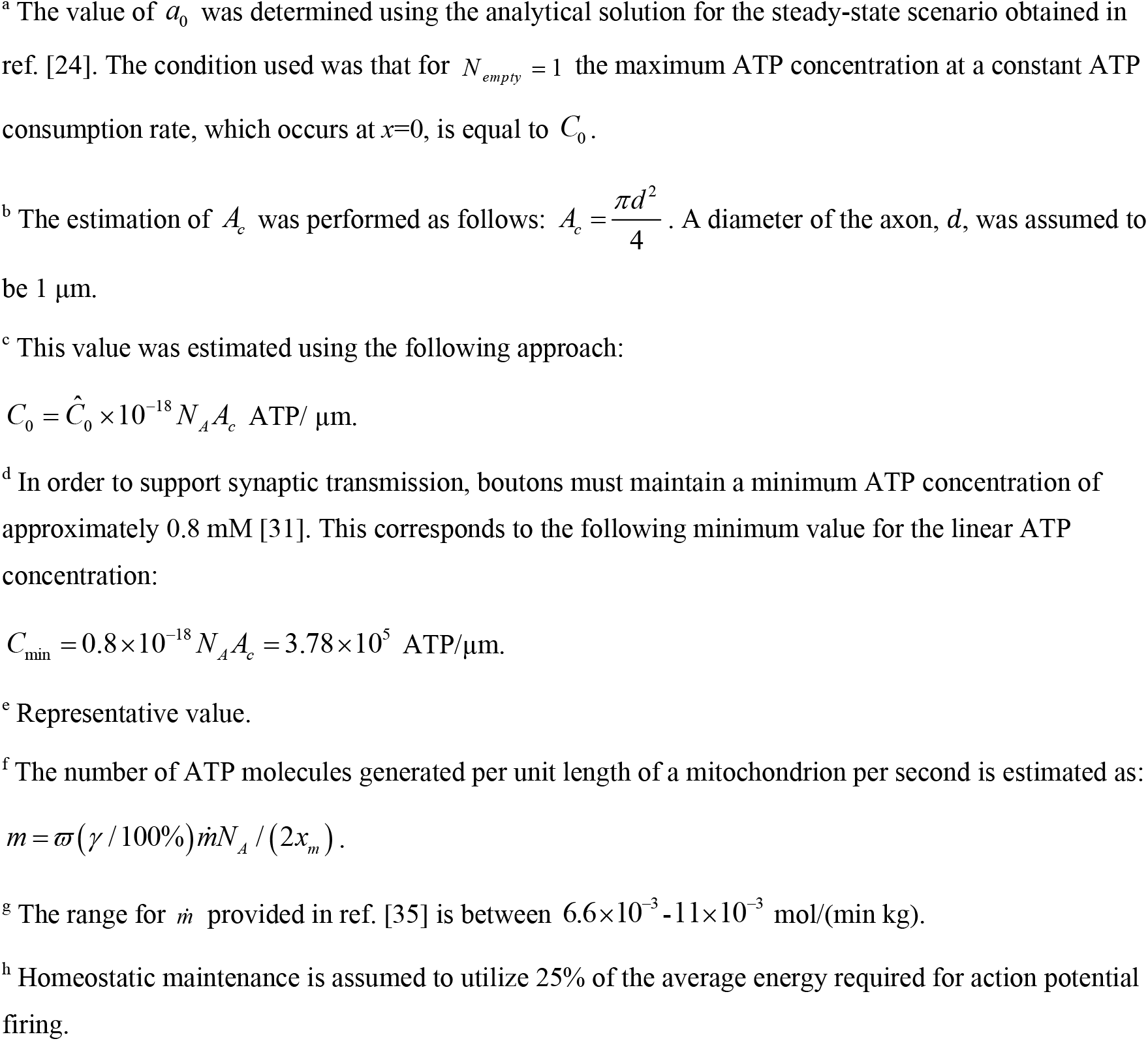
Parameters characterizing ATP production, transport, and consumption in the axon.

| Symbol | Definition | Units | Value or range | Reference(s) | Value used in computations |
| --- | --- | --- | --- | --- | --- |
| $a_0$ | Kinetic constant characterizing the rate of ATP consumption in a bouton | $s^{-1}$ | 0.23 <sup>a</sup> | [24] | 0.23 |
| $A_c$ | Axon's cross-sectional area | $\mu m^2$ | 0.79 <sup>b</sup> | | 0.79 |
| $C_0$ | Typical value of linear ATP concentration in the axon | $\mu\text{m}^{-1}$ | $1.42 \times 10^6$ <sup>c</sup> | [31] | $1.42 \times 10^6$ |
| $C_{\min}$ | Minimum linear ATP concentration required to sustain synaptic transmission | $\mu\text{m}^{-1}$ | $3.78 \times 10^5$ <sup>d</sup> | [31] | $3.78 \times 10^5$ |
| $duty$ | Duty cycle, the percentage of the signal period during which the square wave remains positive | % | | | 40 |
| $D$ | ATP diffusivity in the cytosol | $\mu\text{m}^2 \text{ s}^{-1}$ | 710 – 720<br>in water,<br>~300 in the cytosol | [32,33] | 300 |
| $f$ | Frequency at which neuron fires during the active phase | Hz | 1-200 | [34] | 1 |
| $i_{\text{active}}$ | Number of action potentials that occur during the active phase (5 for the case displayed in Fig. 2) | | | | 5 <sup>e</sup> |
| $i_{\text{total}}$ | Total number of action potentials that propagate down the axon during the active phase plus the number of action potentials that were missed during the quiescent phase (8 for the case displayed in Fig. 2) | | | | 8 <sup>e</sup> |
| $L$ | Average distance between boutons | $\mu\text{m}$ | 3 | [11] | 3, 10 |
| $m$ | ATP production rate per unit length of a mitochondrion | $\mu\text{m}^{-1}\text{s}^{-1}$ | $1.85 \times 10^5$ <sup>f</sup> | [35-37] | $1.29 \times 10^6$ |
| $\dot{m}$ | ATP production rate per unit mass of tissue | $\text{mol s}^{-1} \text{kg}^{-1}$ | $1.1 \times 10^{-4}$<br>$-1.83 \times 10^{-4}$ <sup>g</sup> | [35] | $1.33 \times 10^{-4}$ |
| $N_{\text{empty}}$ | Number of boutons (or half-boutons) in the CV that lack stationary mitochondria. These boutons are supplied with ATP by a bouton containing a stationary mitochondrion | | 1-5 | | 1-5 |
| $2x_m$ | Typical length of a mitochondrion | $\mu\text{m}$ | 0.5-10 | Fig. 2G of ref. [11], ref. [38] | 1 |
| $\gamma$ | Percentage of tissue volume occupied by mitochondria | % | 2-10 | [37] | 5 |
| $\delta$ | Half the width of an axonal varicosity | $\mu\text{m}$ | 1 | Fig. 2E of ref. [11], ref. [39] | 1 |
| $\varpi$ | Mass of an individual mitochondrion | kg | $322 \times 10^{-15}$ | [36] | $322 \times 10^{-15}$ |
| $\Lambda_{\text{homeostatic}}$ | Portion of energy expended on homeostatic maintenance of a bouton | | | | $0.25$ <sup>h</sup> |

<sup>a</sup> The value of $a_0$ was determined using the analytical solution for the steady-state scenario obtained in ref. [24]. The condition used was that for $N_{empty} = 1$ the maximum ATP concentration at a constant ATP consumption rate, which occurs at $x=0$ , is equal to $C_0$ .
<sup>b</sup> The estimation of $A_c$ was performed as follows: $A_c = \frac{\pi d^2}{4}$ . A diameter of the axon, $d$ , was assumed to be 1 $\mu\text{m}$ .
<sup>c</sup> This value was estimated using the following approach:
<sup>d</sup> In order to support synaptic transmission, boutons must maintain a minimum ATP concentration of approximately 0.8 mM [31]. This corresponds to the following minimum value for the linear ATP concentration:
<sup>e</sup> Representative value.
<sup>f</sup> The number of ATP molecules generated per unit length of a mitochondrion per second is estimated as:
<sup>g</sup> The range for $\dot{m}$ provided in ref. [35] is between $6.6 \times 10^{-3}$ - $11 \times 10^{-3}$ mol/(min kg).
<sup>h</sup> Homeostatic maintenance is assumed to utilize 25% of the average energy required for action potential firing.

### 2.1. Equations governing ATP concentration in a CV, which includes one bouton containing a stationary mitochondrion and another bouton lacking a stationary mitochondrion (*N*_*empty*_ = 1)

In the scenario depicted in Fig. 1a, which corresponds to *N*_*empty*_ = 1, a single stationary mitochondrion supplies ATP to the bouton it occupies and to one empty bouton, with half situated to the left of the mitochondrion’s symmetry line at *x*=0 and the other half to the right. The width of an axonal varicosity is denoted by 2 *δ*, and the average distance between consecutive varicosities is *L*. Stationary mitochondria are assumed to be located at every alternate varicosity, resulting in a spacing of 2 *L* + 4*δ* between the centers of two neighboring mitochondria. The conservation of ATP in the segment of a bouton containing a mitochondrion is expressed by the following equation:

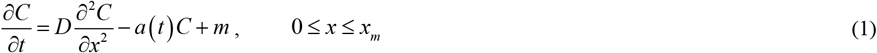

Here, *t* represents time, *x* denotes the linear position along the axon, *D* is the diffusivity of ATP in the cytosol, *m* is the ATP production rate per unit length of the mitochondrion, and *a* characterizes the rate of ATP consumption in a bouton. The second term on the right-hand side of Eq. (1) is based on the assumption that ATP consumption is directly proportional to the concentration of ATP available for binding.

Within a part of varicosity that lacks a stationary mitochondrion, as well as within an empty bouton (Fig. 1a), the ATP conservation equation is expressed as:

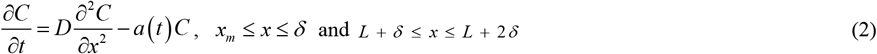

In the intervaricosity spacing (Fig. 1a) ATP conservation is governed by the following equation:

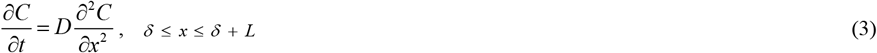

The function *a* (*t*) is modeled based on the following biological considerations. A neuron transmits electrical signals by allowing sodium and potassium ions to flow sequentially into the cell along its axon. While sodium and potassium channels operate based on passive ion flow [26], their function depends on the sodium/potassium pump [27], which works continuously to restore ion gradients and maintain a higher concentration of sodium ions outside the neuron and a higher concentration of potassium ions inside the neuron, enabling the propagation of action potentials. The sodium/potassium pump utilizes the energy of ATP hydrolysis [28,29]. The activity rate of the Na^+^/K^+^ pump is influenced by ion concentrations, which fluctuate after spikes, causing the pump’s energy demands to vary over time. Neurons also require ATP for the release, reuptake, and cycling of neurotransmitters during synchronous neurotransmitter release and synaptic vesicle recycling in presynaptic boutons [28,30].

A basal level of ATP consumption is required in synapses to support essential homeostatic functions [30]. Homeostatic maintenance is assumed to account for Λ_*homeostatic*_ portion of the average energy spent on action potential firing (Fig. 2 is plotted for Λ_*homeostatic*_ = 0.25, which corresponds to 25%). Additionally, it is assumed that *i*_*active*_ action potentials are followed by a quiescent phase lasting the equivalent duration of *i*_*total*_ − *i*_*active*_ action potentials. This behavior is represented by the following periodic rate of ATP consumption in a bouton:

**Fig. 2.**
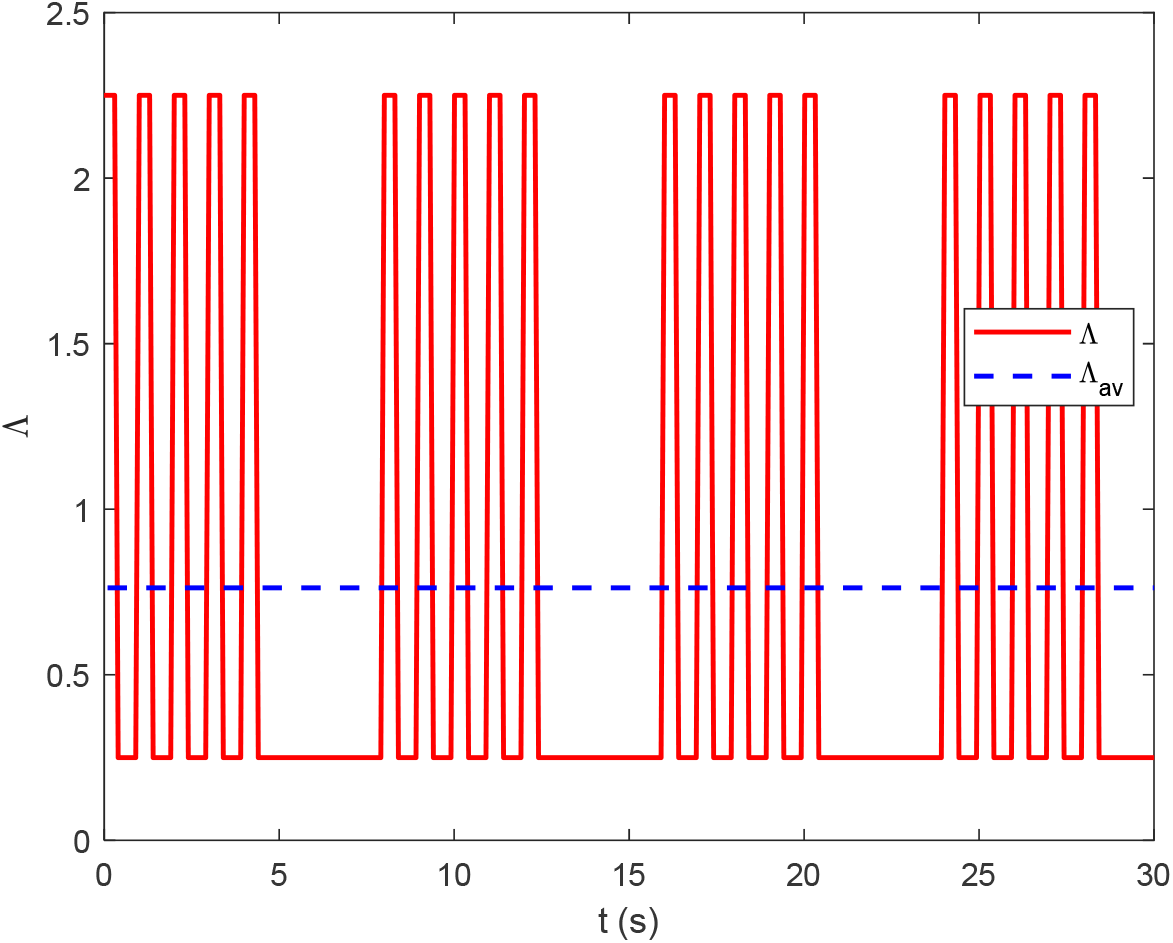
The time-dependent factor, Λ, of the ATP consumption rate for the scenario of oscillating ATP demand, simulating a neuron firing a series of five spikes followed by a quiescent period. A constant basal ATP consumption rate is assumed to be necessary for maintaining metabolic equilibrium. Additionally, each spike involves energy costs associated with spike generation, neurotransmitter release, reuptake, and synaptic vesicle reloading. The time-averaged value of Λ is also displayed.

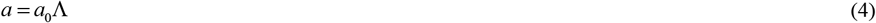

where

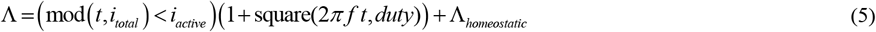

In Eq. (5), *i*_*active*_ is the number of action potentials generated during the active phase (Fig. 2 is plotted for *i*_*active*_ = 5), *i*_*total*_ is the number of action potentials generated during the active phase plus those omitted during the quiescent phase (Fig. 2 is plotted for *i*_*total*_ = 8). Also, in Eq. (5), mod (*t,i*_*total*_) denotes the remainder after the division of time *t* by *i*_*total*_; (mod (*t,i*_*total*_) < *i*_*active*_) equals 1 if mod (*t,i*_*total*_) < *i*_*active*_, otherwise, it equals 0. Additionally, square(2*π f t, duty*) is a periodic step function with a period of 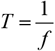, where *f* represents the frequency of neuronal firing during active phases and *duty* is the duty cycle that indicates the percentage of the signal’s phase during which the square wave is positive. If *duty* = 50%, the function square resembles the sine function but produces a square wave with values of – 1 and 1. Fig. 2 is plotted for *duty*=40%.

To facilitate a comparison between the scenario with an oscillating ATP consumption rate, as described by Eqs. (4) and (5) and illustrated in Fig. 2, and the scenario with constant ATP consumption, the average ATP consumption rate was calculated as follows:

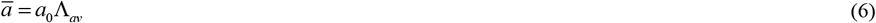

where

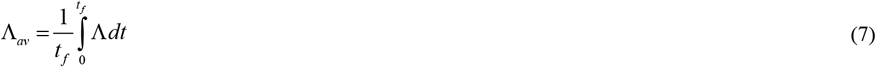

In the computations presented in this paper, a value of *t*_*f*_= 100 s was used, yielding an average ATP consumption rate of

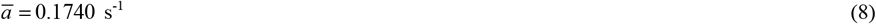

Given the periodic nature of the axon segment illustrated in Fig. 1a, symmetry conditions are applied at the segment’s endpoints, *x* = 0 and *x* = *L* + 2*δ*. Additionally, the continuity of ATP concentration and ATP flux is enforced at all internal interfaces within the segment. Accordingly, the boundary conditions for Eqs. (1)-(3) are as follows:

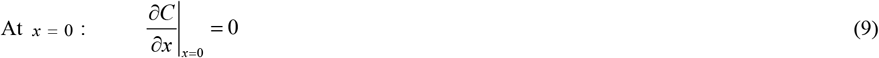

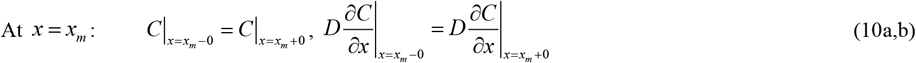

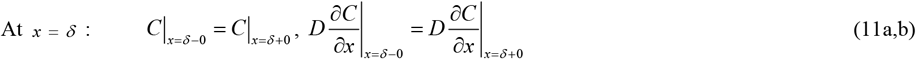

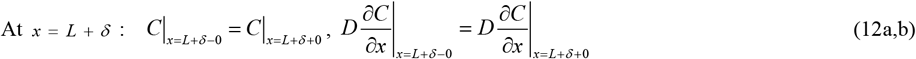

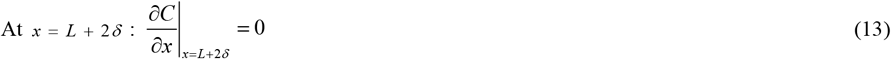

Healthy axons function under homeostatic conditions for decades, making the ATP concentration at long times biologically relevant. Consequently, the initial condition is not critical. In this study, a zero initial condition was utilized:

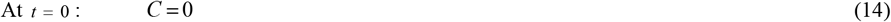

### 2.2. Equations governing ATP concentration in a CV containing one bouton with a stationary mitochondrion and multiple boutons without stationary mitochondria (*N*_*empty*_ > 1)

In the scenario shown in Fig. 1b (*N*_*empty*_ = 5), a single mitochondrion supports 9 empty boutons, with 4.5 to the left and 4.5 to the right of the bouton containing a mitochondrion. The CV displayed in Fig. 1b includes only half a bouton containing a mitochondrion and 4.5 boutons without mitochondria. This situation is described by the following equations:

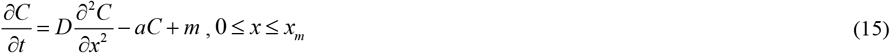

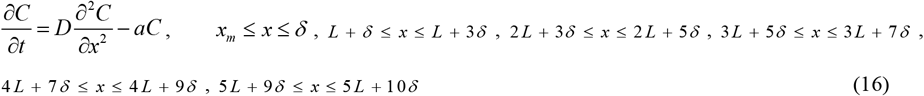

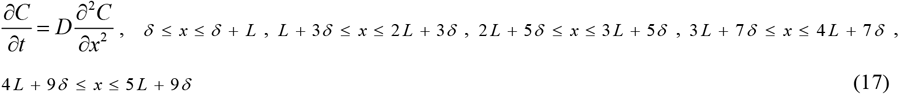

At the interfaces corresponding to *x* = *x*_*m*_, *x* = *δ*, *x* = *L* + *δ*, *x* = *L* + 3*δ*, *x* = 2*L* + 3*δ*, *x* = 2*L* + 5*δ*, *x* = 3*L* + 5*δ*, *x* = 3*L* + 7*δ*, *x* = 4*L* + 7*δ*, *x* = 4*L* + 9*δ*, and *x* = 5*L* + 9*δ* the ATP concentration and the ATP diffusion flux are assumed to remain continuous. At *x* = 0 and *x* = 5*L* +10*δ* a symmetry condition, 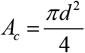, is applied, reflecting the assumption that identical CVs exist on both sides of the computational domain.

### 2.3. Simplified equation governing ATP concentration under the lumped capacitance approximation

Assuming that the ATP concentration is independent of position within the CV (i.e., independent of *x*), which is a reasonable approximation for large ATP diffusivity, the balance between ATP consumption and production at steady-state can be expressed as follows:

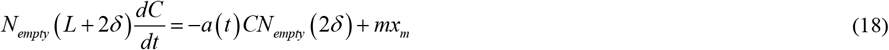

Eq. (18), along with the initial condition given in Eq. (14), can be numerically solved to determine *C* (*t*).

For the case of constant ATP consumption in boutons, i.e. *a* = *a*, the analytical solution of Eq. (18), utilizing the initial condition specified in Eq. (14), is given by:

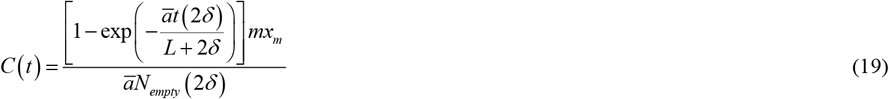

For *t* →∞, Eq. (19) simplifies to:

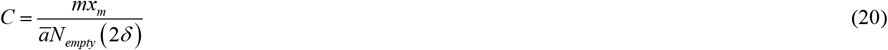

The criterion for the validity of applying the lumped capacitance approximation can be established as follows. Consider the scenario depicted in Fig. 1a. The ATP flux from a bouton containing a mitochondrion to one without a mitochondrion can be estimated by analyzing the ATP consumption within the bouton lacking a mitochondrion:

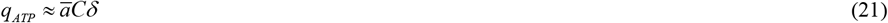

On the other hand, the ATP flux due to diffusion from a bouton containing a mitochondrion to a bouton lacking a mitochondrion can be estimated as follows:

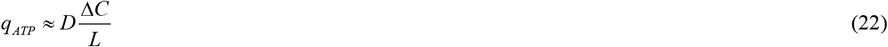

where Δ*C* represents the difference in ATP concentration between a bouton containing a mitochondrion and one lacking a mitochondrion.

By equating the right-hand sides of Eqs. (20) and (21) and solving for Δ*C* / *C*, the following criterion is obtained:

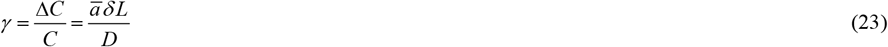

The lumped capacitance approximation is valid when *γ* ≪ 1. Using the parameter values from Table 1 and Eq. (8), the value of *γ* was estimated to be 0.0017. This parameter is analogous to the Biot number in convection-conduction problems.

### 2.4. Numerical solution

To solve the system of partial differential equations (PDEs) governing ATP concentration variation along the axon, a single-domain approach was employed [40]. This method enables the solution of a single equation across the entire computational domain, with changes in the equation between subdomains accounted for by varying the coefficients in different subdomains. Consequently, the equation adapts to the appropriate form within each subdomain. For instance, for the computational domain illustrated in Fig. 1b and described in Section 2.2, the ATP conservation equation for the entire computational domain is as follows:

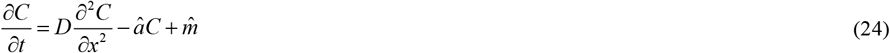

Within the region occupied by a mitochondrion, the coefficients in Eq. (24) are defined as:

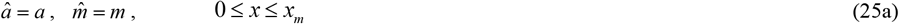

In the portion of the bouton not occupied by a mitochondrion, as well as in boutons lacking mitochondria, the coefficients are:

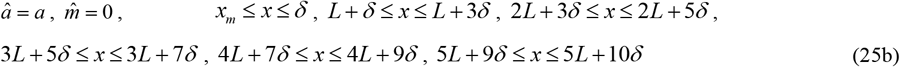

In the intervaricosity spacing, the coefficients are defined as:

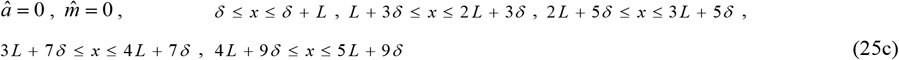

Using the single-domain approach ensures that the numerical solution inherently maintains the continuity of both ATP concentration and ATP flux at the interfaces between subdomains, *x* = *x*_*m*_, *x* = *δ*, *x* = *L* + *δ, x* = *L* + 3*δ, x* = 2 *L* + 3*δ*, *x* = 2 *L* + 5*δ, x* = 3 *L* + 5*δ*, *x* = 3 *L* + 7 *δ*, *x* = 4 *L* + 7 *δ*, *x* = 4 *L* + 9*δ*, and *x* = 5 *L* + 9*δ*.

Boundary conditions are required only at the external boundaries of the domain:

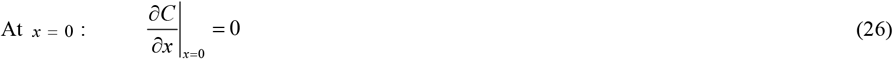

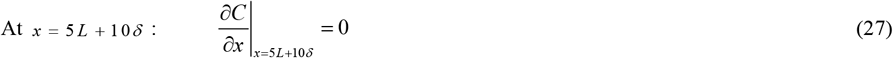

Eq. (24), along with the boundary conditions specified in Eqs. (26) and (27) and the initial condition specified in Eq. (14), was numerically solved using a well-validated MATLAB’s PDEPE solver (MATLAB R2023a, MathWorks, Natick, MA, USA). The solution was computed on a (1000, 50000) mesh in the (*x,t*) domain. The computation, performed on a PC equipped with an Intel Core i9 processor, required approximately 10 minutes to complete. A mesh-independence study was conducted using a finer (2000, 100000) mesh within the (*x,t*) domain.

Eq. (18) obtained under the lumped capacitance approximation, along with the initial condition given by Eq. (14), was numerically solved using MATLAB’s ODE23 solver (MATLAB R2020b, MathWorks, Natick, MA, USA). To achieve high accuracy, the solver’s relative and absolute tolerances (RelTol and AbsTol) were set to 1e-10.

## 3. Results

### 3.1. Exploring the effect of spontaneous firing on ATP levels in boutons with and without mitochondria

For the case of oscillating ATP consumption, a scenario is simulated in which the neuron fires a series of five spikes followed by a quiescent period during which it skips three spikes (Fig. 2). It is assumed that the energy required for homeostatic maintenance constitutes 25% of the average energy cost of spike firing.

The ATP concentration increases rapidly during quiescent periods and decreases five consecutive times in response to each square wave of ATP consumption thereafter (Fig. 3). This indicates that the propagation of neuronal action potentials can lead to a reduction in ATP concentration, with successive axonal potentials causing a stepwise decline, where each step corresponds to an individual action potential. These findings align with those reported in ref. [14], which suggested that spontaneous neuronal firing may act as a “release valve,” responding to stalled ATP production during periods of low energy demand. Stalled ATP production poses a significant risk to neurons, as it can result in the release of highly harmful ROS from mitochondria [14].

**Fig. 3.**
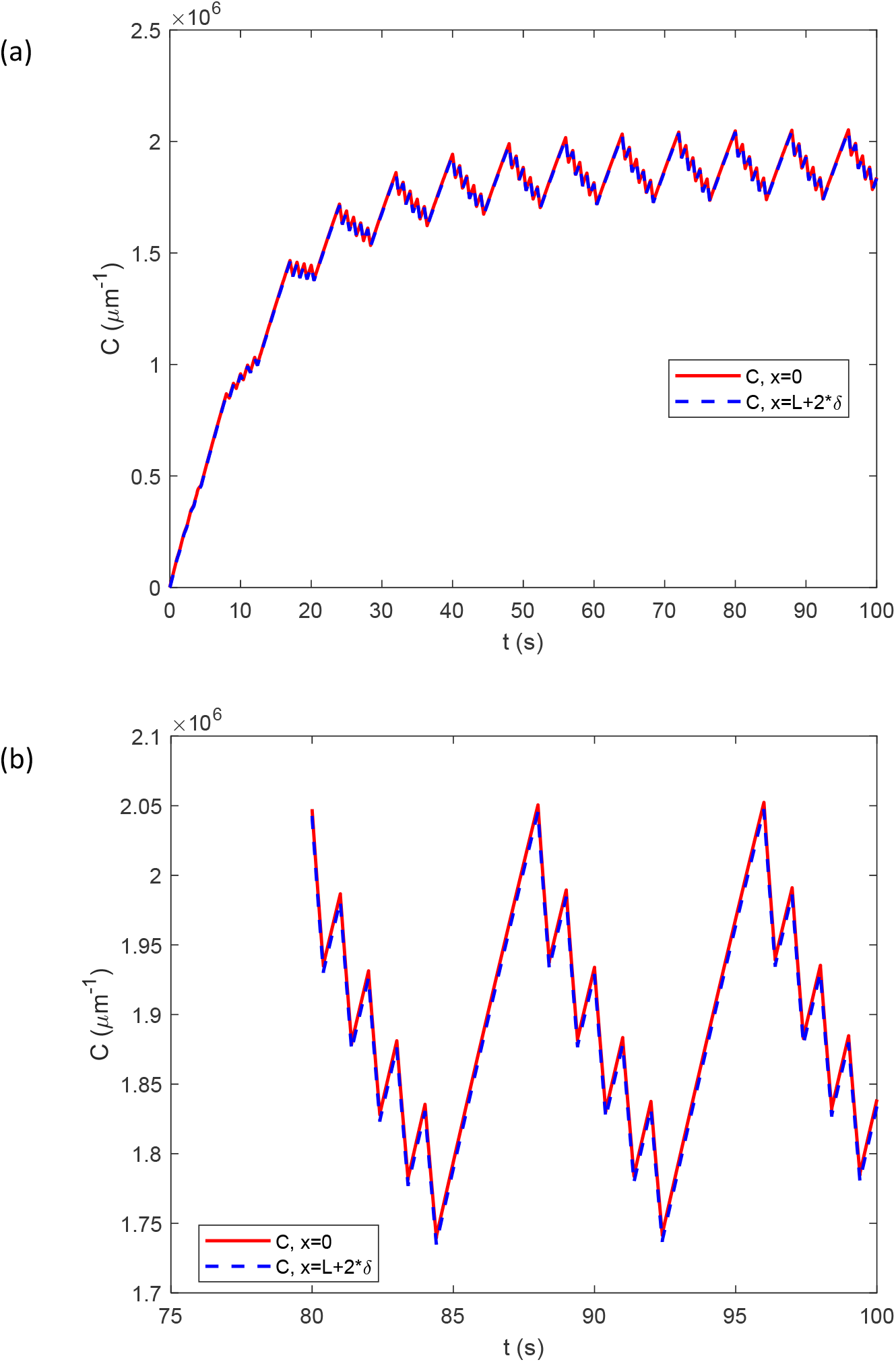
The case of oscillating ATP consumption displayed in Fig. 2. ATP concentration vs time for the scenario where a stationary mitochondrion is present at every alternate varicosity (depicted in Fig. 1a, *N*_*empty*_ = 1). (a) ATP concentration at *x*=0 (the center of a bouton with a stationary mitochondrion) and at *x* = *L* + 2*δ* (the center of a bouton without a stationary mitochondrion). (b) A magnified section of the graph in Fig. 3a.

### 3.2. Assessing ATP support from a single donated mitochondrion to empty boutons

In the case of constant ATP consumption at steady state, increasing the number of empty boutons surrounding a bouton containing a mitochondrion does not result in a significant reduction in ATP concentration with increasing distance from the mitochondrion (Fig. 4a). This demonstrates that ATP diffusivity is sufficiently high to transport ATP from the bouton containing the mitochondrion to the end of the CV, passing through multiple empty boutons, without a notable decline in ATP concentration along the *x-*axis. This conclusion is supported by the results (indicated by markers) obtained using the lumped capacitance approximation, which assumes infinitely high ATP diffusivity and thus disregards any variation in ATP concentration along the *x*-axis. The analytical solution derived using the lumped capacitance approximation is provided in Eq. (20).

**Fig. 4.**
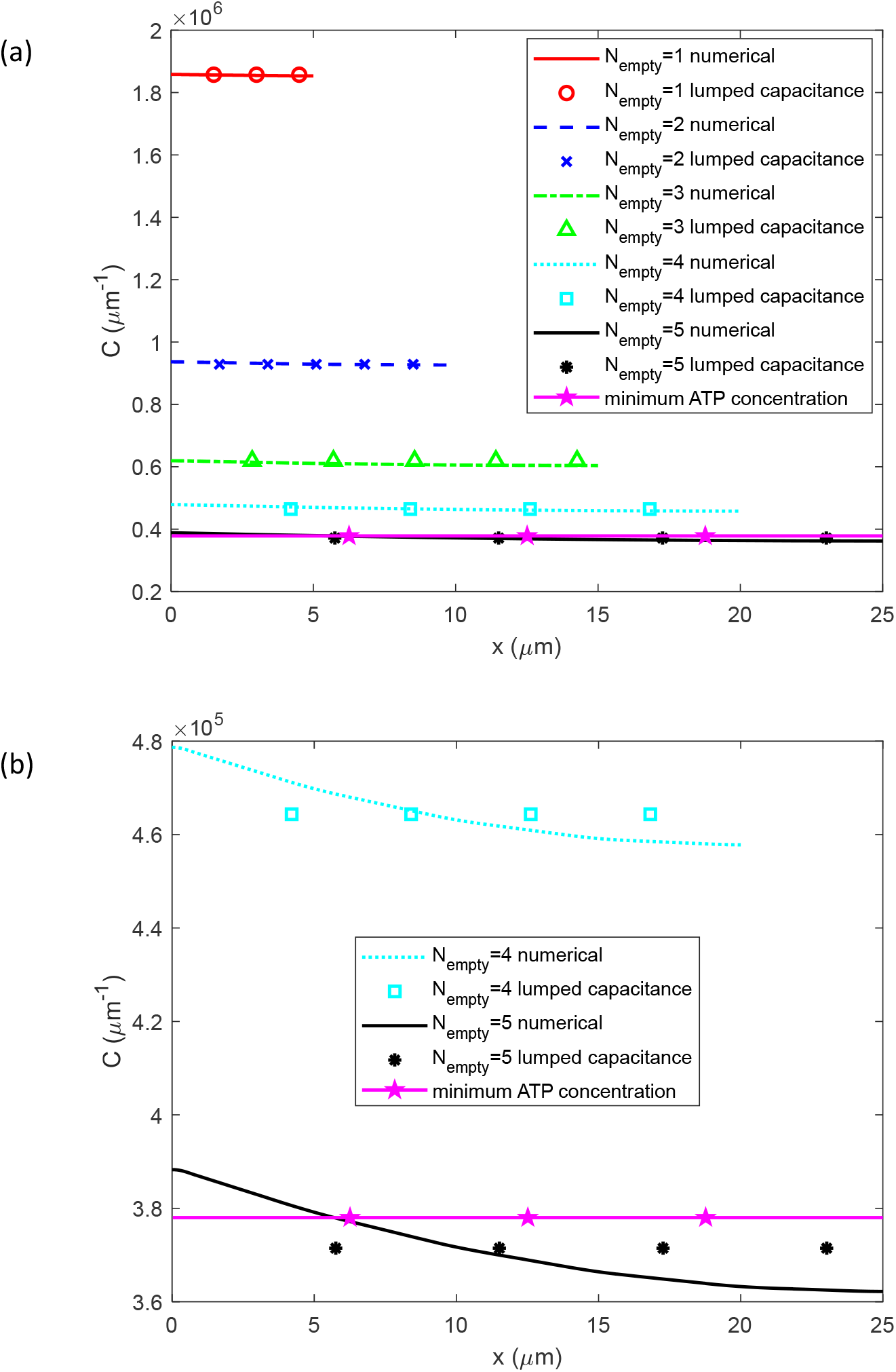
The scenario with constant ATP consumption, *a* = *ā*. (a) The ATP concentration is plotted vs the distance from the center of a bouton containing a mitochondrion, considering scenarios with varying numbers (1 to 5) of empty boutons. The minimum ATP concentration required to support synaptic transmission is also plotted. (b) A magnified section of the graph in Fig. 4a for four and five empty boutons showing the variation of numerically computed ATP concentration with *x*.

As the number of empty boutons increases, the total ATP consumption within the CV rises. This occurs because all boutons—including the one containing a mitochondrion and the empty (lacking stationary mitochondria) boutons—are actively firing, which consumes energy. Consequently, the average ATP concentration decreases as the number of empty boutons increases. For the case of five empty boutons, the ATP concentration drops to the threshold required to support synaptic transmission (Fig. 4a).

A magnified section of the graph in Fig. 4a for four and five empty boutons reveals that, for a physiologically relevant ATP diffusivity of 300 µm^2^/s (see Table 1), the ATP concentration varies with *x* (Fig. 4b). The maximum ATP concentration is found at the center of the bouton containing a mitochondrion, while the minimum occurs at the center of the farthest empty bouton. This variation is not apparent in Fig. 4a due to the larger range of the *y*-axis.

When ATP consumption in an axonal segment with multiple empty boutons oscillates due to neuron firing, the ATP concentration decreases after each spike but recovers to its initial level by the end of the quiescent period (Fig. 5). As the number of empty boutons increases, the average ATP concentration decreases. Notably, when the number of empty boutons reaches five, the ATP concentration may fall below the minimum level required to sustain synaptic transmission (Fig. 5).

**Fig. 5.**
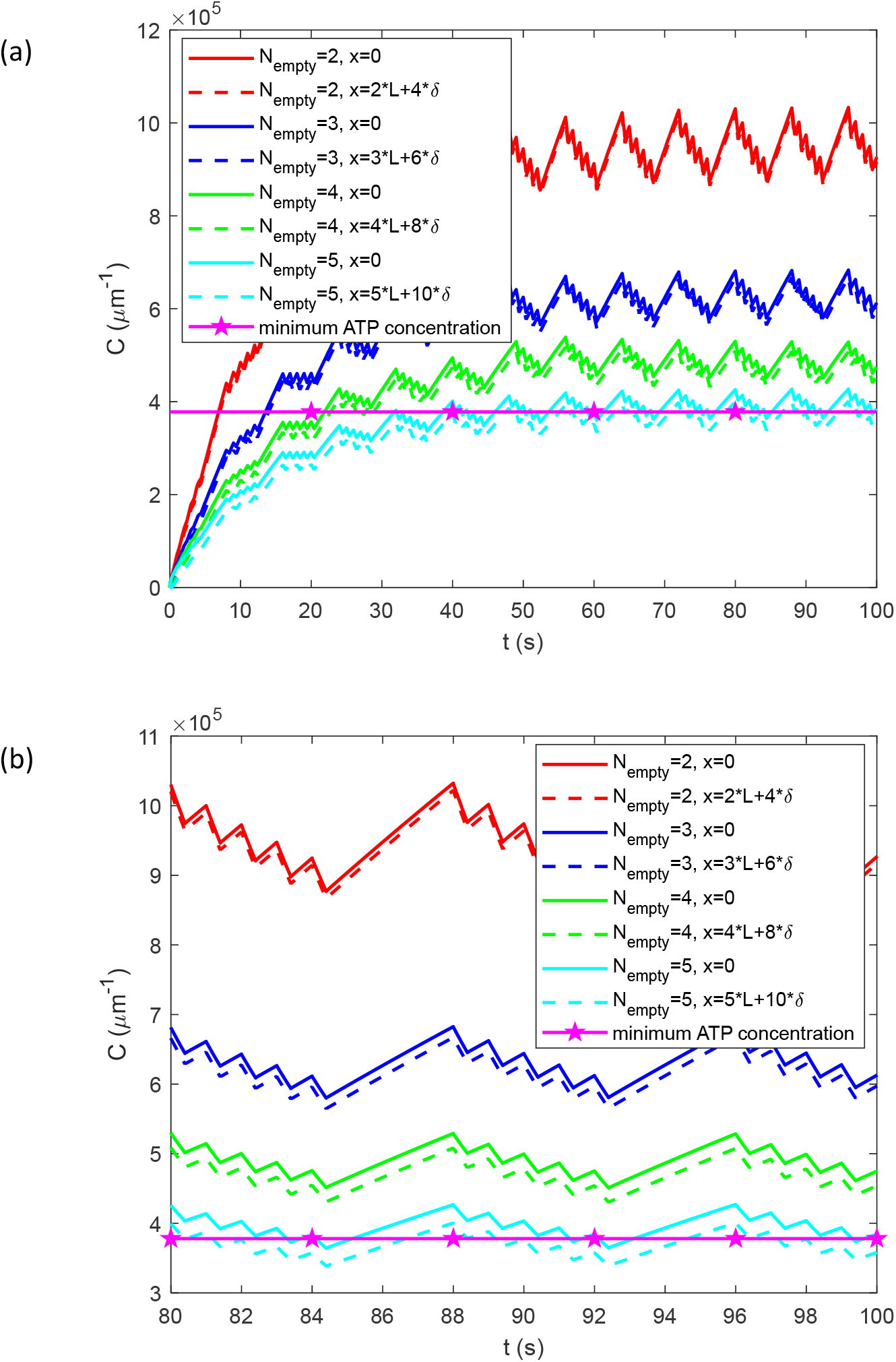
The case of oscillating ATP consumption displayed in Fig. 2. ATP concentration vs time for the scenario where a stationary mitochondrion is present in a bouton surrounded by several empty boutons (*N*_*empty*_ = 2, 3, 4, 5). (a) ATP concentration at *x*=0 (the center of a bouton with a stationary mitochondrion) and in the center of the last empty bouton in the CV (the case for *N*_*empty*_ = 5 is depicted is in Fig. 1b). (b) A magnified section of the graph in Fig. 5a.

### 3.3. Assessing the validity of the lumped capacitance approximation

Fig. 6 demonstrates strong consistency between the ATP concentrations for the case with constant ATP consumption derived from solving the complete set of partial differential equations, Eqs. (1)–(3) and those obtained using the lumped capacitance approximation for the transient scenario (given by Eq. (19)). The transient solutions converge to the lumped capacitance solution derived for the steady-state scenario (Eq. (20)) as time progresses (Fig. 6). Fig. 7 demonstrates good agreement between the ATP concentration calculated by solving the full set of partial differential equations, Eqs. (1)–(3), and the simplified equation derived using the lumped capacitance approximation, Eq. (18).

**Fig. 6.**
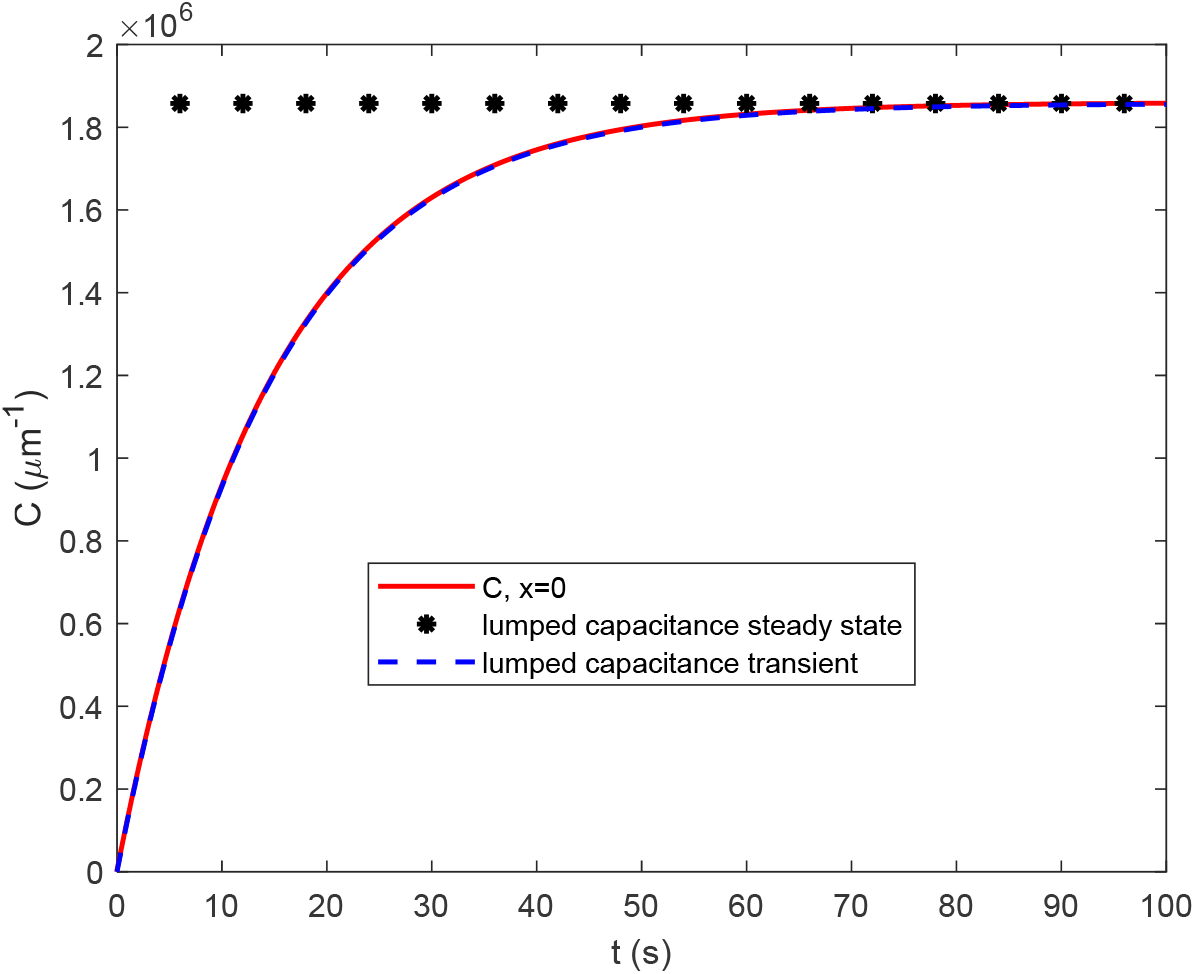
The case of constant ATP consumption. ATP concentration vs time for the scenario where a stationary mitochondrion is present at every alternate varicosity (depicted in Fig. 1a, *N*_*empty*_ = 1). The ATP concentration at *x*=0 (the center of a bouton containing a stationary mitochondrion) computed by solving the full set of PDEs compared with the ATP concentrations calculated using the lumped capacitance approximation for both steady-state and transient scenarios.

**Fig. 7.**
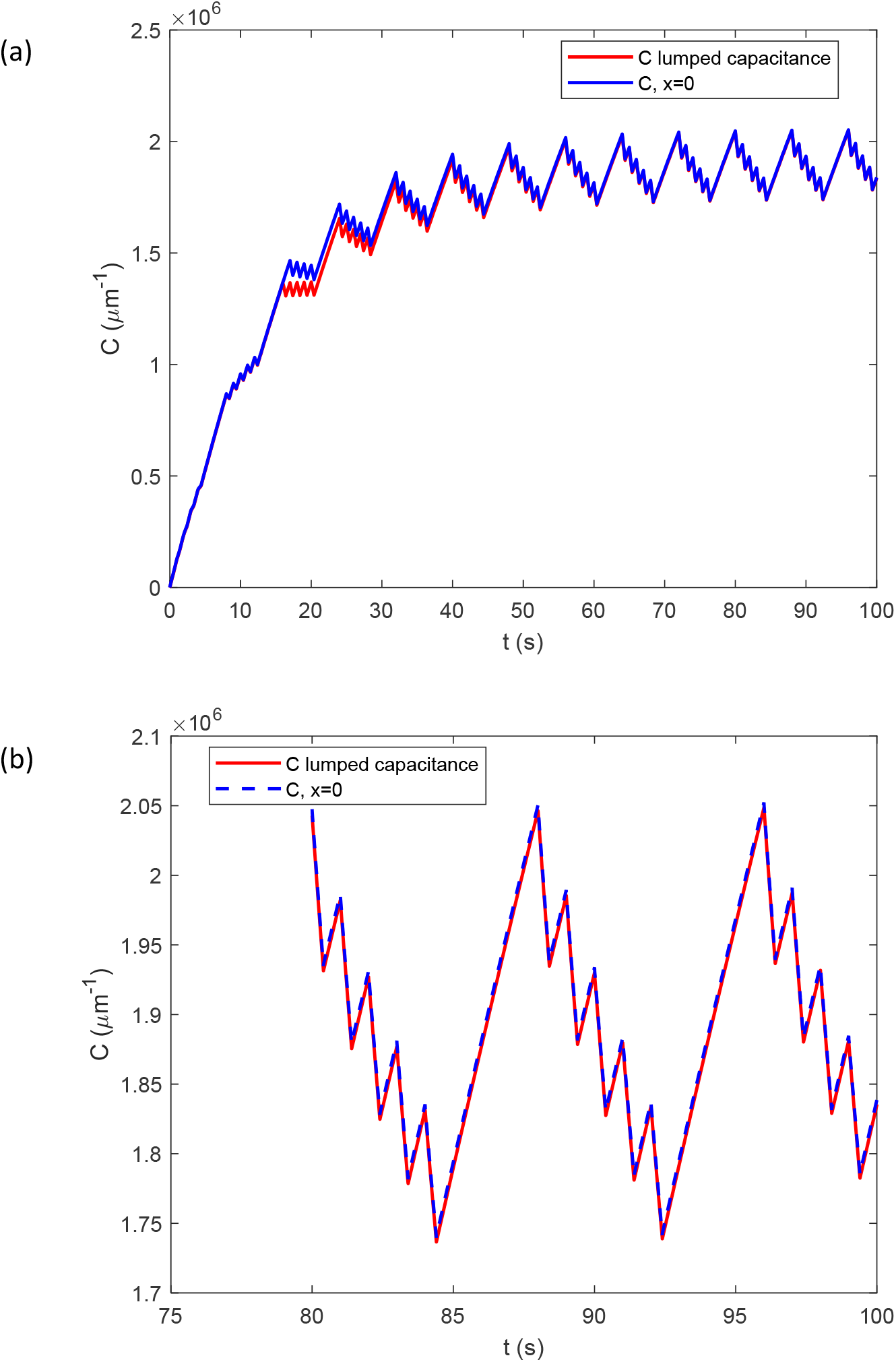
The case of oscillating ATP consumption displayed in Fig. 2. ATP concentration vs time for the scenario where a stationary mitochondrion is present at every alternate varicosity (depicted in Fig. 1a, *N*_*empty*_ = 1). (a) ATP concentration computed utilizing the lumped capacitance approximation and ATP concentration at *x*=0 (the center of a bouton with a stationary mitochondrion) computed solving the full set of PDEs. (b) A magnified section of the graph in Fig. 7a.

The results presented in Fig. 7 appear counterintuitive at first glance. One might expect the spikes shown in Fig. 3 to arise from ATP diffusion. However, the findings in Fig. 7 suggest that due to the high diffusivity of ATP, its concentration remains nearly uniform along the *x*-axis. Instead, the spikes are primarily driven by ATP production in mitochondria and ATP consumption in boutons. Consequently, ATP concentration varies predominantly with time rather than spatial position.

## 4. Conclusions

Spontaneous neuronal firing plays a protective role in preventing the release of ROS from mitochondria during periods of low energy demand, when mitochondrial ADP levels become insufficient at mitochondrial complex V, as noted in ref. [14]. Simulation results reveal that consecutive axonal potentials cause a stepwise decrease in the ATP concentration, with each drop corresponding to an individual action potential. This reduction in ATP concentration is linked to its conversion into ADP, resulting in an increased ADP concentration, which prevents dangerous conditions at complex V.

Mitochondria can be transferred to degenerating neurons via tunneling nanotubes (TNTs), fragile intercellular structures that facilitate such transmission. Since the number of mitochondria transferred through TNTs is expected to be low [21,22], it is crucial to assess how many neighboring boutons lacking mitochondria a single bouton containing a donated mitochondrion can support. Simulations show that under constant ATP consumption, the average ATP concentration declines as the number of empty boutons increases. When five boutons lack mitochondria, the ATP concentration drops to the minimum threshold required for synaptic transmission. Notably, this drop in ATP concentration is not confined to the most distal empty bouton but occurs throughout the entire axonal segment, a phenomenon attributed to the high diffusivity of ATP. In scenarios where ATP consumption fluctuates due to neuronal firing, the presence of five empty boutons can also cause ATP levels to fall below the critical threshold, impairing synaptic activity.

A criterion *γ*, defined by Eq. (23), is established. It demonstrates that when *γ* ≪ 1, variations in ATP concentration along the axonal length can be neglected, allowing the problem to be simplified using the lumped capacitance approximation, which assumes that ATP concentration is solely time-dependent. The parameter *γ* is analogous to the Biot number in convection-conduction problems.

## Abbreviations

ADP: adenosine diphosphate
ATP: adenosine triphosphate
CV: control volume
PDE: partial differential equation
ROS: reactive oxygen species

## CRediT authorship contribution statement

AVK is the sole author of this paper.

## Acknowledgment

The author acknowledges the support provided by the National Science Foundation (grant CBET- 2042834) and the Alexander von Humboldt Foundation through the Humboldt Research Award.

